# Inter-regional interactions uncouple within-region inhibition stabilization and paradoxical responses

**DOI:** 10.64898/2026.09.09.750483

**Authors:** Yue Kris Wu, Kenneth D. Miller

## Abstract

Paradoxical responses, in which excitatory perturbations of inhibitory neurons reduce their firing rates, are widely regarded as a hallmark of inhibition-stabilized networks (ISNs) and have been observed in multiple brain regions. However, because brain regions are interconnected by long-range projections, it remains unclear whether a paradoxical response observed in a given region reflects inhibition stabilization within that region or instead arises from distributed interactions among regions. Here, using analytically tractable multi-region population models and numerical simulations, we show that inter-regional connections can dissociate local inhibition stabilization from paradoxical responses. The condition for a paradoxical response in a given region is jointly determined by recurrent excitation within that region, feedback mediated through other regions, and the dynamics of those other regions. Consequently, inter-regional coupling can generate a paradoxical response in a region that is not locally inhibition-stabilized or abolish it in a region that is. Thus, in an interconnected network, a paradoxical response is neither necessary nor sufficient for local inhibition stabilization. These findings demonstrate that local perturbation responses cannot generally be interpreted solely in terms of local circuit dynamics and highlight the importance of accounting for inter-regional connections when inferring the local circuit properties of individual brain regions.

## Introduction

Recurrent excitation is a widespread feature of neural circuits and supports neural computations such as input amplification (Murphy and Miller, 2009; Peron et al., 2020), pattern completion (Hopfield, 1982; Guzman et al., 2016), active filtering (Histed, 2025; Deveau et al., 2026), and working memory (Amit and Brunel, 1997; Wong and Wang, 2006; Hilty and Miller, 2026). When sufficiently strong, however, recurrent excitation can render the excitatory subnetwork unstable unless it is dynamically stabilized by inhibitory feedback. Networks operating in this regime are known as inhibition-stabilized networks (ISNs), in which inhibition stabilizes an otherwise unstable excitatory subnetwork (Tsodyks et al., 1997; Ozeki et al., 2009; Sadeh and Clopath, 2021).

A widely recognized hallmark of an ISN is the *paradoxical inhibitory response*, in which increasing the excitatory input to inhibitory neurons decreases their steady-state firing rates, whereas decreasing the input increases them (Tsodyks et al., 1997; Ozeki et al., 2009). This counterintuitive response arises because, following a positive perturbation to the inhibitory population, inhibitory activity initially increases and suppresses the excitatory population, thereby reducing the recurrent excitatory drive to inhibitory neurons. When this indirect reduction in recurrent drive outweighs the direct effect of the perturbation, inhibitory activity decreases at steady state.

Paradoxical responses have been observed in multiple brain regions and are commonly interpreted as evidence that the perturbed local circuit operates in an inhibition-stabilized regime (Li et al., 2019; Sanzeni et al., 2020; de Jong et al., 2023). This interpretation, however, is largely based on theoretical results derived for single-region circuit models (Tsodyks et al., 1997; Ozeki et al., 2009; Sanzeni et al., 2020; Wu and Gjorgjieva, 2023). Local circuits in the brain are instead embedded within networks of regions interconnected by abundant long-range projections (Markov et al., 2014). A local perturbation applied to one region can alter activity in other regions, and the resulting activity changes can then feed back onto the perturbed circuit (Guo et al., 2017; Javadzadeh et al., 2024; Clark and Beiran, 2025; Mastrogiuseppe et al., 2026). These interactions create inter-regional feedback loops that are absent from isolated-circuit models. It therefore remains unclear whether a paradoxical response observed in a given region reflects inhibition stabilization within that region or can instead be generated by distributed interactions among regions.

Here, we address this question through mathematical analysis of multi-region population models and numerical simulations. We show that, in the presence of inter-regional connections, a paradoxical response within a given region is jointly determined by recurrent excitation within that region, feedback mediated through other regions, and the dynamics of those other regions. Consequently, inter-regional connections can generate a paradoxical response in a region that is not locally inhibition-stabilized and abolish it in a region that is. Thus, in an interconnected network, a paradoxical response is neither a necessary nor a sufficient indicator of local inhibition stabilization. We further decompose inter-regional feedback into effective excitatory and inhibitory pathways, providing a mechanistic account of how long-range connectivity can promote or oppose paradoxical responses. Together, these findings highlight the importance of considering inter-regional connections when inferring the local circuit properties of a brain region from its perturbation responses.

## Results

To determine how inter-regional connections affect the relationship between local inhibition stabilization and paradoxical inhibitory responses, we consider a population model of *N* brain regions, each comprising one excitatory (E) population and one inhibitory (I) population. The firing-rate dynamics of the network are governed by

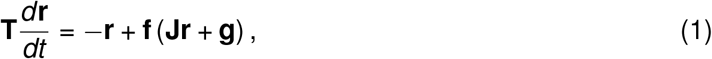

where

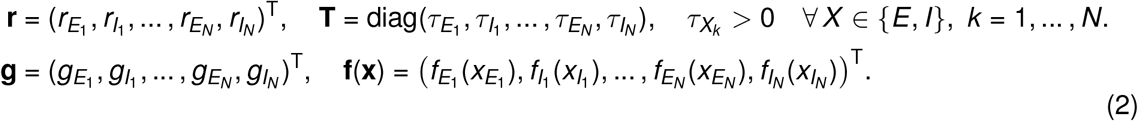

Here, **r** and **g** denote the firing rate and external input vectors, respectively, and **T** is the diagonal matrix of the corresponding firing-rate time constants. The matrix **J** specifies local and inter-regional connectivity. The activation function **f** acts element-wise on the total input, with each component assumed to be continuously differentiable and monotonically increasing.

The connectivity matrix can be partitioned into 2 × 2 blocks:

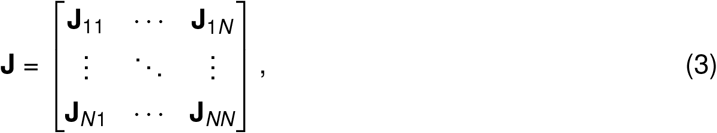

where **J**_*kℓ*_ represents connections from region *ℓ* to region *k* . Both local (*k* = *ℓ*) and inter-regional (*k*≠ *ℓ*) blocks have the form

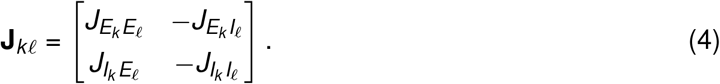

Each 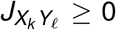 denotes the magnitude of the connection from population *Y* in region *ℓ* to population *X* in region *k*, with *X, Y* ∈ {*E, I*}. The explicit minus signs indicate connections originating from inhibitory populations.

To characterize local inhibition stabilization and paradoxical responses, we analyze the linearized dynamics around a fixed point. We assume the network is initially at a fixed point **r**^∗^ satisfying

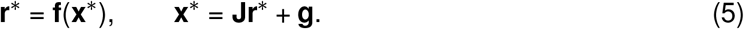

We consider a perturbation *δ***g** to the external input and denote the corresponding deviation from the fixed-point firing rates by *δ***r** = **r** − **r**^∗^. Assuming that the perturbation is sufficiently small, we approximate the dynamics to first order in *δ***g** and *δ***r**. The linearized dynamics are

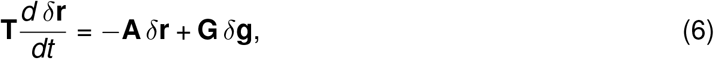

where

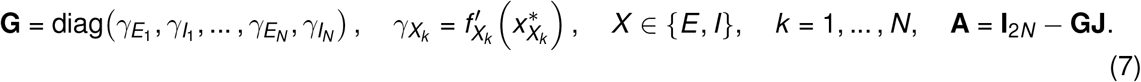

Here, **I**_2*N*_ denotes the 2*N* × 2*N* identity matrix. We assume that all gains 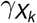 are strictly positive at the fixed point.

### Local inhibition stabilization

Region *p* is locally inhibition-stabilized if its excitatory population *E*_*p*_ is unstable when all other populations are held fixed, while its local E–I circuit is stable when populations in all other regions are held fixed (Ozeki et al., 2009). To assess the intrinsic stability of *E*_*p*_, we hold *I*_*p*_ and all populations in the other regions fixed at their fixed-point values. The single eigenvalue of the Jacobian of the *E*_*p*_ subsystem is

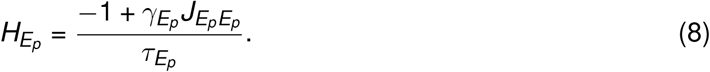

Since 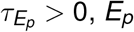 is unstable precisely when

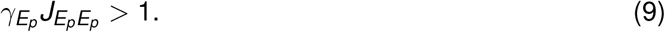

To assess the stability of the local E–I circuit, we allow both *E*_*p*_ and *I*_*p*_ to vary while holding all populations in other regions fixed. The local E–I Jacobian is

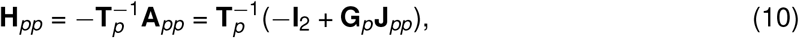

where 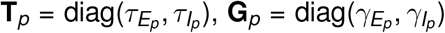, and **A**_*pp*_ = **I**_2_ − **G**_*p*_**J**_*pp*_. The local E–I circuit is stable when

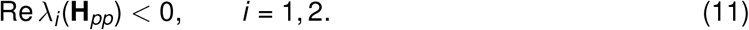

Region *p* is therefore locally inhibition-stabilized when both Equations (9) and (11) hold.

Additionally, we require the full interconnected network to be stable. The Jacobian of the full network is

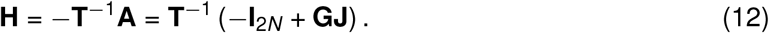

The full network is asymptotically stable when all eigenvalues of **H** have negative real parts:

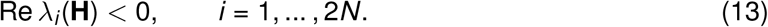

### Paradoxical responses

We next summarize the general condition for a paradoxical response of the inhibitory population *I*_*p*_, following Miller and Palmigiano (2020). For a small constant input perturbation, setting the time derivative in Equation (6) to zero gives

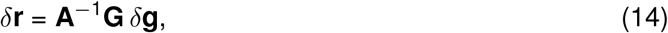

assuming that **A** is nonsingular.

For a perturbation applied selectively to *I*_*p*_,

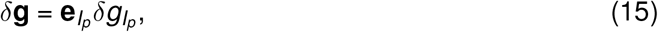

where 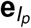 is the coordinate vector associated with *I*_*p*_. The resulting change in the steady-state activity of *I*_*p*_ is

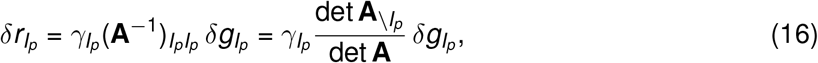

where 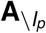 denotes the principal submatrix obtained by deleting the row and column associated with *I*_*p*_.

A paradoxical response occurs when the inhibitory activity changes in the direction opposite to the applied perturbation:

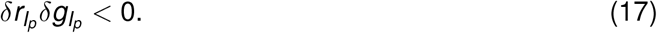

Because 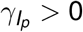, the above condition is equivalent to

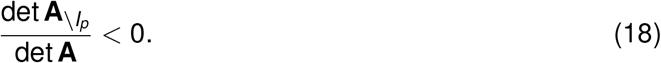

Under the full-network stability condition in Equation (13), all eigenvalues of **H** have negative real parts, and therefore det(−**H**) *>* 0. Since **A** = −**TH** and det **T** *>* 0, we obtain det **A** = det **T** det(−**H**) *>* 0.

Thus, the inhibitory response is paradoxical precisely when

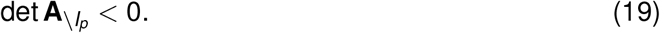

As shown in Miller and Palmigiano (2020), the condition det 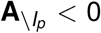 implies that the Jacobian of the complementary subsystem obtained by excluding *I*_*p*_ has a positive determinant and that this subsystem has an odd number of unstable modes. Here, unstable modes are defined as eigen-vectors associated with eigenvalues having positive real parts, with the corresponding eigenvalues counted according to their algebraic multiplicities.

### Local inhibition stabilization and paradoxical responses in an isolated region

Let *R* denote all excitatory and inhibitory populations outside region *p*. To recover the standard single-region result, we disconnect region *p* from *R* by setting

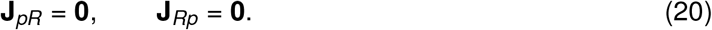

Unlike holding other regions fixed at their baseline activities, removing inter-regional connections eliminates their baseline input and may therefore shift the operating point. Quantities evaluated at the fixed point of the isolated configuration are denoted by hats. These differ in general from the unhatted quantities used to assess local inhibition stabilization at the interconnected fixed point. In the isolated configuration, the excitatory population *E*_*p*_ is unstable when *I*_*p*_ is held fixed if

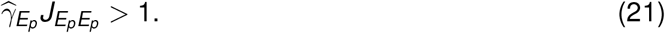

For a perturbation applied selectively to *I*_*p*_, the resulting change in its steady-state activity in the isolated E–I circuit is

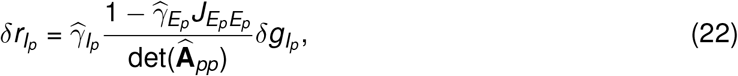

where 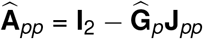. If the isolated E–I circuit is asymptotically stable, then 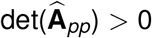. Because 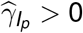, the inhibitory response is paradoxical precisely when the local excitatory population is unstable with inhibition held fixed:

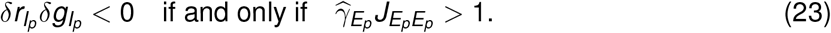

Thus, in a stable isolated E–I circuit, local inhibition stabilization is equivalent to a paradoxical response of the inhibitory population, recovering the standard single-region result (Tsodyks et al., 1997; Ozeki et al., 2009).

### Inter-regional coupling dissociates local inhibition stabilization from paradoxical responses

We next examine how inter-regional coupling modifies the relationship between local inhibition stabilization and paradoxical responses. We return to the fixed point of the interconnected network and denote the associated quantities without hats. Because the paradoxical-response condition is determined by det 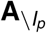, we partition the matrix 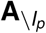 into *E*_*p*_ and *R* components:

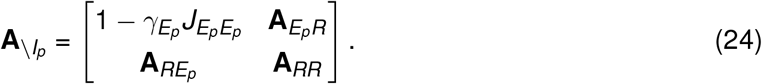

Assuming that **A**_*RR*_ is nonsingular, since

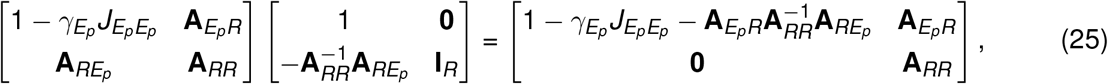

we have

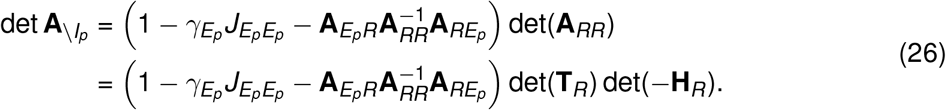

Defining the feedback mediated through *R* as

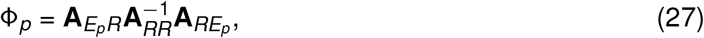

we obtain

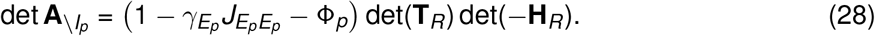

The term 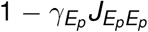 describes the intrinsic stability of the local excitatory population, whereas Φ_*p*_ represents feedback that leaves *E*_*p*_, propagates through *R*, and returns to *E*_*p*_. The factor det(**T**_*R*_) det(−**H**_*R*_) captures the contribution of the dynamics within *R*. Thus, the paradoxical-response condition in the interconnected network is jointly determined by local recurrent excitation, feedback mediated through other regions, and the dynamics of those other regions.

The resulting change in the steady-state activity of *I*_*p*_ induced by a perturbation applied selectively to *I*_*p*_ can then be written as

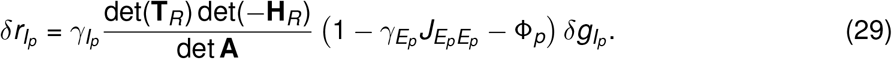

For a stable full network, det **A** *>* 0. Because 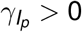 and det(**T**_*R*_) *>* 0, the response is paradoxical under the following conditions:

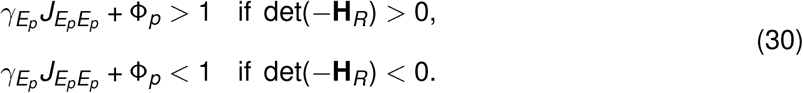

#### Paradoxical responses without local inhibition stabilization

If 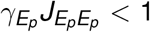, region *p* is not locally inhibition-stabilized. Nevertheless, *I*_*p*_ responds paradoxically when 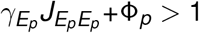 if det(−**H**_*R*_) *>* 0, or when the reverse inequality holds if det(−**H**_*R*_) *<* 0.

#### Local inhibition stabilization without a paradoxical response

If 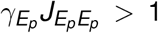 and the local E–I circuit is stable when populations in all other regions are held fixed, region *p* is locally inhibition-stabilized. Nevertheless, its response is nonparadoxical when 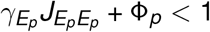 if det(−**H**_*R*_) *>* 0, or when the reverse inequality holds if det(−**H**_*R*_) *<* 0.

Thus, inter-regional coupling can generate a paradoxical response in a region that is not locally inhibition-stabilized or abolish the response in a region that is. Consequently, a paradoxical response is neither necessary nor sufficient to establish local inhibition stabilization.

### Pathway decomposition of inter-regional feedback

Having established the general conditions under which inter-regional coupling dissociates local inhibition stabilization from paradoxical responses, we next interpret Φ_*p*_ as feedback mediated by directed connectivity pathways through *R*. Using 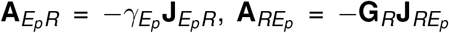, and **A**_*RR*_ = **I**_*R*_ − **G**_*R*_**J**_*RR*_, we obtain

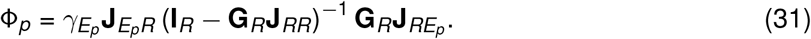

When the spectral radius *ρ*(**G**_*R*_**J**_*RR*_) *<* 1, the inverse admits the convergent Neumann-series expansion

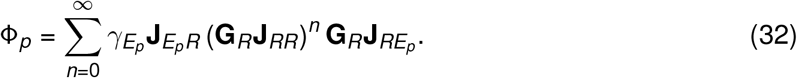

The term indexed by *n* sums pathways that leave *E*_*p*_, undergo *n* internal transitions within *R*, and return to *E*_*p*_.

Consider a pathway *π* : *E*_*p*_ −→ *X*_1_ −→ · · · −→ *X*_*ℓ*_ −→ *E*_*p*_, where *X*_1_, …, *X*_*ℓ*_ are populations in *R*. Its contribution to Φ_*p*_ is

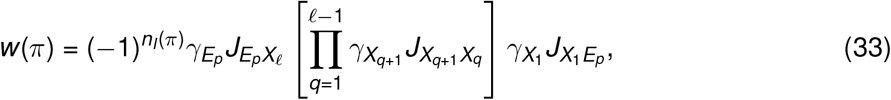

where *n*_*I*_(*π*) denotes the number of inhibitory connections along the pathway. For *ℓ* = 1, the product is empty and is defined to equal one. Each connection originating from an inhibitory population contributes one minus sign, including the return from *X*_*ℓ*_ to *E*_*p*_ when *X*_*ℓ*_ is inhibitory.

A pathway with even *n*_*I*_(*π*) contributes positive feedback and is termed an effective excitatory pathway, whereas a pathway with odd *n*_*I*_(*π*) contributes negative feedback and is termed an effective inhibitory pathway. To sum the excitatory and inhibitory contributions separately, we impose the stronger sufficient condition *ρ*(|**G**_*R*_**J**_*RR*_|) *<* 1, where the absolute value is taken entrywise. Under this condition, the pathway expansion converges absolutely, and the sums of the contributions from the effective excitatory and inhibitory pathways are therefore finite. We define the corresponding effective excitatory and inhibitory feedback strengths as

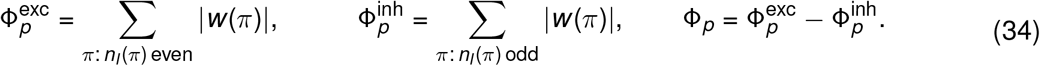

This absolute-convergence condition also implies *ρ*(**G**_*R*_**J**_*RR*_) *<* 1 and hence det(−**H**_*R*_) *>* 0. The paradoxical-response condition therefore becomes

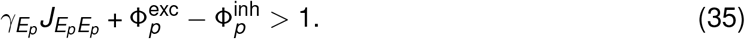

Thus, under the absolute-convergence condition, effective excitatory feedback lowers the amount of local recurrent excitation required for a paradoxical response, whereas effective inhibitory feed-back raises it.

When 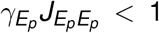, region *p* is not locally inhibition-stabilized, but sufficiently strong effective excitatory feedback through *R* can generate a paradoxical response. Mechanistically, a positive perturbation to *I*_*p*_ initially increases inhibition and suppresses *E*_*p*_. Effective excitatory pathways reinforce this suppression, further reducing excitatory drive to *I*_*p*_. If the indirect network effect outweighs the direct perturbation, the steady-state activity of *I*_*p*_ decreases, producing a paradoxical response despite the absence of local inhibition stabilization.

Conversely, when region *p* is locally inhibition-stabilized 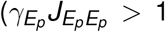 and the local E–I circuit is stable), sufficiently strong effective inhibitory feedback through *R* can abolish a paradoxical response. The effective inhibitory pathways counteract the suppression of *E*_*p*_, limiting the resulting loss of excitatory drive to *I*_*p*_. If the direct perturbation outweighs this reduction, inhibitory activity increases at steady state, producing a nonparadoxical response despite local inhibition stabilization.

### Simulations illustrate the dissociation induced by inter-regional coupling

To illustrate the theoretical results, we simulated a two-region network with threshold-linear activation functions. The simulation parameters were chosen such that the activities of all populations remained nonzero throughout the simulations. For each connectivity configuration, we adjusted the external inputs to maintain the same baseline firing rate of 5 in all populations, allowing comparisons across configurations at a fixed operating point (see Numerical Simulations). We first configured both region 1 and region 2 as non-ISNs (Figure 1A). Accordingly, in each region, the eigenvalue of the Jacobian of the excitatory subsystem had a negative real part (Figure 1B), while the full E–I circuit of each isolated region was asymptotically stable, with all Jacobian eigenvalues having negative real parts (Figure 1C). In the uncoupled configuration, an excitatory perturbation applied selectively to *I*_1_ increased its steady-state activity and therefore did not elicit a paradoxical response (Figure 1D).

**Fig. 1.**
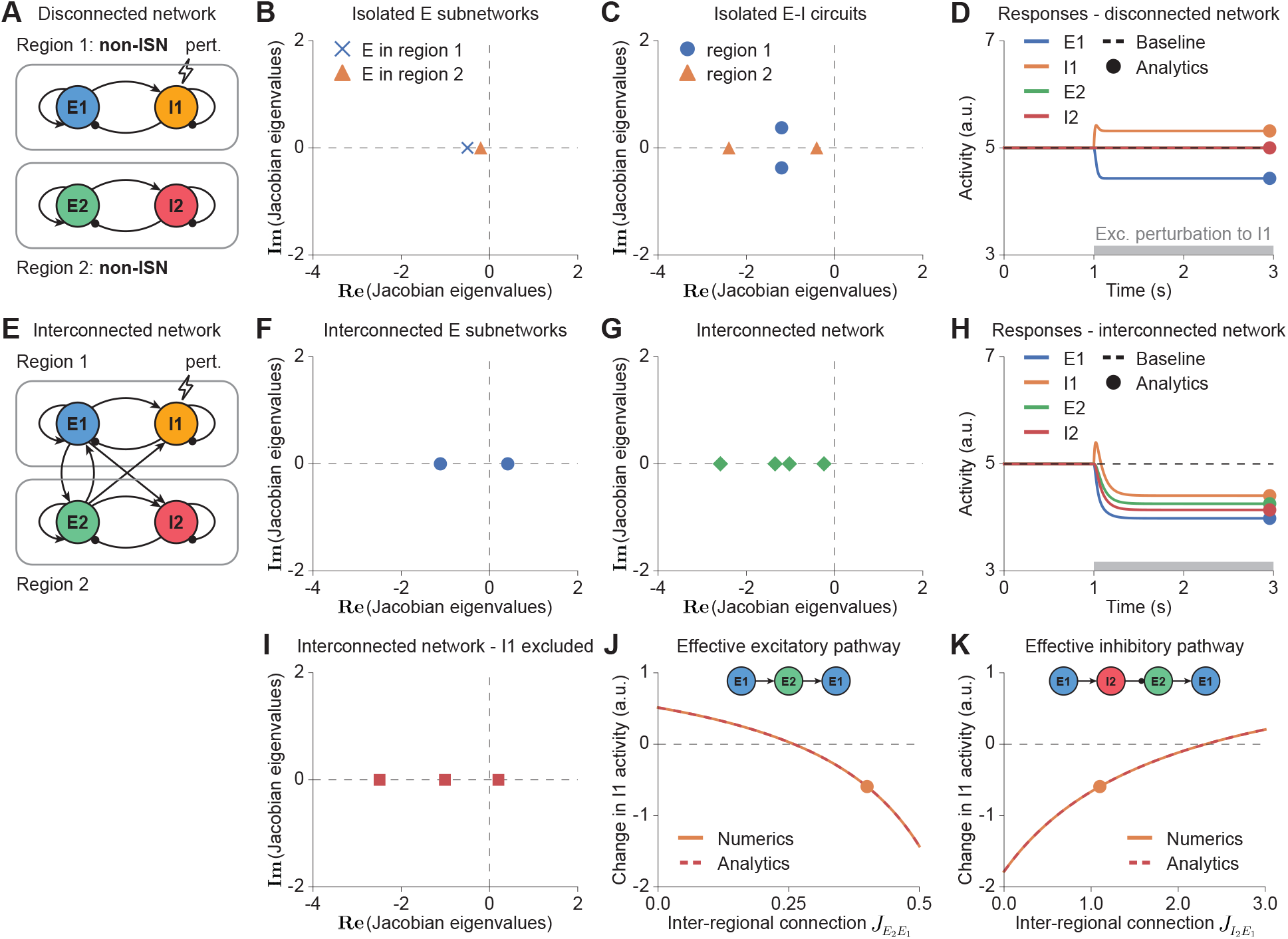
Inter-regional connections can induce a paradoxical inhibitory response in a non-ISN coupled to a non-ISN. A–D. Disconnected configuration. **A**. Schematic of two disconnected regions, each comprising an excitatory (E) population and an inhibitory (I) population. Both regions are non-ISNs in isolation. **B**. Eigenvalues of the Jacobians of the isolated E subnetworks. The eigenvalues are scaled by the common time constant *τ* and are therefore dimension-less. The eigenvalues of the E subnetwork of region 1 (cross) and of region 2 (triangle) both have negative real parts, indicating that each E subnetwork is stable by itself and thus neither isolated region is inhibition-stabilized. **C**. Eigenval-ues of the Jacobians of the isolated E–I circuits. The real parts of all eigenvalues of both regions are negative, indicating that both regions are stable in isolation. Together, **B** and **C** establish that both regions are non-ISNs. **D**. Responses to an excitatory perturbation applied to the inhibitory population in region 1 (*I*_1_) beginning at 1 s (gray bar). Dashed lines indicate baseline activities, and dots denote the analytical post-perturbation steady-state activities. *I*_1_ activity increases in response to the perturbation, indicating the absence of a paradoxical inhibitory response. **E–K**. Interconnected configuration. **E–H**. Similar to **A–D**, but for the interconnected network. The coupled excitatory subnetwork now has one unstable mode, but the full interconnected network is stable. In the presence of inter-regional connections, when an excitatory perturbation is applied to *I*_1_, its steady-state activity decreases, *i*.*e*., its response is paradoxical, despite region 1 being a non-ISN in isolation. **I**. Eigenvalues of the Jacobian of the interconnected network with *I*_1_ excluded. The reduced network without *I*_1_ has an odd number of unstable eigenmodes (one in this case), consistent with the paradoxical response of *I*_1_ in **H. J**. Effect of the inter-regional connection *J*_*E*2*E*1_, which contributes to positive inter-regional feedback, on the perturbation-induced change in *I*_1_ activity (*δr*_*I*1_). Solid and dashed lines indicate numerical and analytical results, respectively. The dot marks the example shown in **E–I**. The unperturbed region (region 2) has zero (an even number of) unstable eigenmodes, as shown in **C**. Increasing *J*_*E*2*E*1_ decreases the perturbation-induced change in *I*_1_ activity. **K**. Similar to **J** but for the inter-regional connection *J*_*I*2*E*1_, which contributes to negative inter-regional feedback. Increasing *J*_*I*2*E*1_ increases the perturbation-induced change in *I*_1_ activity.

We then coupled the regions through inter-regional excitatory connections, with no inter-regional inhibitory connections included for simplicity (Figure 1E). After coupling, the interconnected excitatory subnetwork became unstable, with an eigenvalue having a positive real part (Figure 1F), while the full interconnected E–I network remained asymptotically stable (Figure 1G).

Because the threshold-linear activation functions have unit slope above threshold, all active populations have unit gain. Assuming that all populations remain above threshold before and after the perturbation therefore gives **G** = **I**. In all examples, the local E–I circuit of the perturbed region, region 1, is stable when the other region is held fixed. Because local connection strengths are un-changed by coupling and all populations retain unit gain, the local Jacobian **H**_11_ is identical in the disconnected and interconnected configurations. Thus, the local E–I stability shown in Figure 1C also holds after coupling, and region 1 is locally inhibition-stabilized precisely when 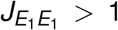. Region 2 is allowed to be stable or unstable when isolated. Under this assumption, the change in the steady-state activity of *I*_1_, induced by a perturbation applied selectively to *I*_1_, can be written as

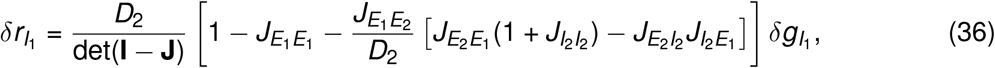

where

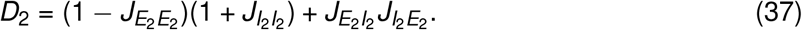

Here, *D*_2_ = det(**I**_2_ − **J**_22_), where **J**_22_ is the local connectivity matrix of region 2. For *D*_2_≠ 0, its sign reflects the parity of the number of unstable modes in the Jacobian of isolated region 2. Specifically, *D*_2_ *>* 0 for an even number of unstable modes and *D*_2_ *<* 0 for an odd number.

After the regions were coupled, the same perturbation decreased the steady-state activity of *I*_1_, producing a paradoxical response even though region 1 was not locally inhibition-stabilized (Figure 1H). Moreover, the Jacobian of the complementary subsystem obtained by excluding *I*_1_ had an odd number of eigenvalues with positive real parts (Figure 1I), consistent with the theoretically established relationship between a population’s paradoxical response and the unstable modes of its complementary subsystem.

For the two-region network, the net inter-regional feedback is

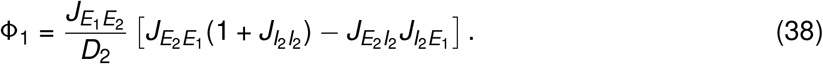

This exact expression accounts for recurrent interactions within region 2 and does not require convergence of the pathway expansion.

In the present example, region 2 is stable in isolation, so *D*_2_ *>* 0. We then write Φ_1_ = Ψ^(+)^ − Ψ^(−)^, where

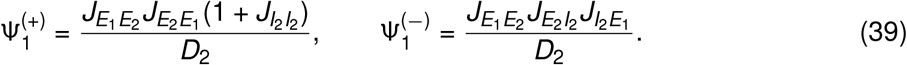

These nonnegative quantities, 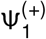 and 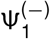, represent the magnitudes of the positive and negative algebraic contributions to Φ_1_, respectively. They are obtained from the exact inverse of the region-2 block and should not be identified individually with the parity-separated pathway sums defined before.

For *D*_2_ *>* 0, the inhibitory response is paradoxical precisely when 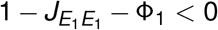, provided the full network remains stable. With all other connections fixed, increasing the strength of the inter-regional connection 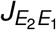, which is involved in the effective excitatory pathway *E*_1_ → *E*_2_ → *E*_1_, increases 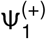 and hence increases Φ_1_, decreasing 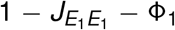. Conversely, increasing the strength of the inter-regional connection 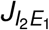, which is involved in the effective inhibitory pathway *E*_1_ → *I*_2_ → *E*_2_ → *E*_1_, increases 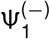 and decreases Φ_1_, increasing 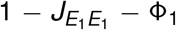. Note that these changes in the inter-regional feedback affect whether the paradoxical-response inequality is satisfied, but they do not alone determine how the response amplitude changes. This is because the positive denominator det(**I** − **J**) generally varies with connection strength as well. In particular, increasing Φ_1_ need not make an already paradoxical response more negative, and decreasing Φ_1_ need not make an already nonparadoxical response more positive. To determine how 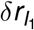 changes with the inter-regional connections 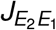 and 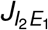, we differentiate it with respect to each connection strength while holding all other connections and the perturbation amplitude fixed:

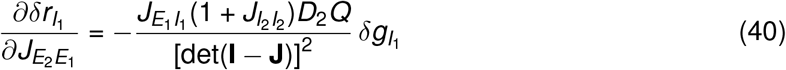

and

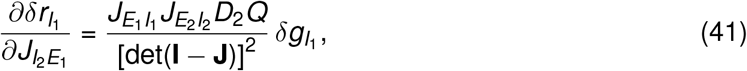

where

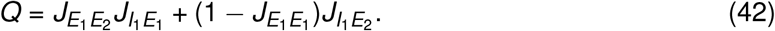

For a positive perturbation and strictly positive connections appearing in the prefactors, the signs of these derivatives are determined by the sign of *D*_2_*Q*. When *D*_2_*Q >* 0, increasing 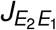 decreases 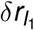, whereas increasing 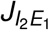 increases it. These trends reverse when *D*_2_*Q <* 0. Note that *Q* is independent of 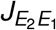 and 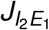. Moreover, because all connection magnitudes are nonnegative, *Q* ≥ 0 whenever 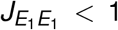. To allow consistent comparisons across conditions, we chose all interconnected parameter sets such that *Q >* 0. In the present example, *D*_2_ *>* 0. Increasing 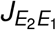 decreased 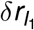, thereby promoting a paradoxical response (Figure 1J), whereas increasing 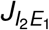 increased 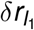, thereby opposing a paradoxical response (Figure 1K).

These simulations show that inter-regional coupling can induce a paradoxical response without local inhibition stabilization. This phenomenon also occurred when the perturbed non-ISN region (region 1) was coupled to an ISN (Figure S1) or to an unstable network with either an even (Figure S2) or an odd (Figure S3) number of unstable eigenmodes. In the latter two cases, although region 2 was unstable in isolation (Figures S2C and S3C), inter-regional coupling rendered the full interconnected network asymptotically stable (Figures S2G and S3G). Because *Q >* 0 for these parameter sets, the signs of the response derivatives were determined by *D*_2_. When region 2 had an odd number of unstable modes (Figure S3C), *D*_2_ *<* 0. Increasing 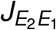 increased 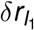 (Figure S3J), whereas increasing 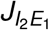 decreased 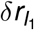 (Figure S3K). These effects were opposite to those observed when region 2 was stable (Figures 1J, K and S1J, K) or had an even number of unstable eigenmodes (Figure S2J, K), for which *D*_2_ *>* 0.

Conversely, when region 1 was locally inhibition-stabilized and exhibited a paradoxical response in isolation, inter-regional connections could abolish that response while preserving the local E–I stability and yielding a stable interconnected network, whether region 2 was an ISN (Figure 2), a non-ISN (Figure S4), or an unstable network with either an even (Figure S5) or an odd (Figure S6) number of unstable modes. All of these parameter sets satisfied *Q >* 0, yielding the same dependence of 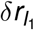 on 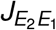 and 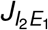 as in the corresponding simulations with a non-ISN region 1. When region 2 was stable or had an even number of unstable modes, *D*_2_ *>* 0. Increasing 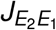 decreased 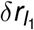 (Figures 2J, S4J, and S5J), whereas increasing 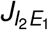 increased 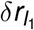 (Figures 2K, S4K, and S5K). When region 2 had an odd number of unstable modes, *D*_2_ *<* 0, and these trends reversed. Increasing 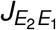 increased 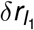 (Figure S6J), whereas increasing 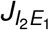 decreased 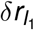 (Figure S6K). Together, these simulations illustrate the theoretical prediction that inter-regional connections can dissociate local inhibition stabilization from paradoxical responses.

**Fig. 2.**
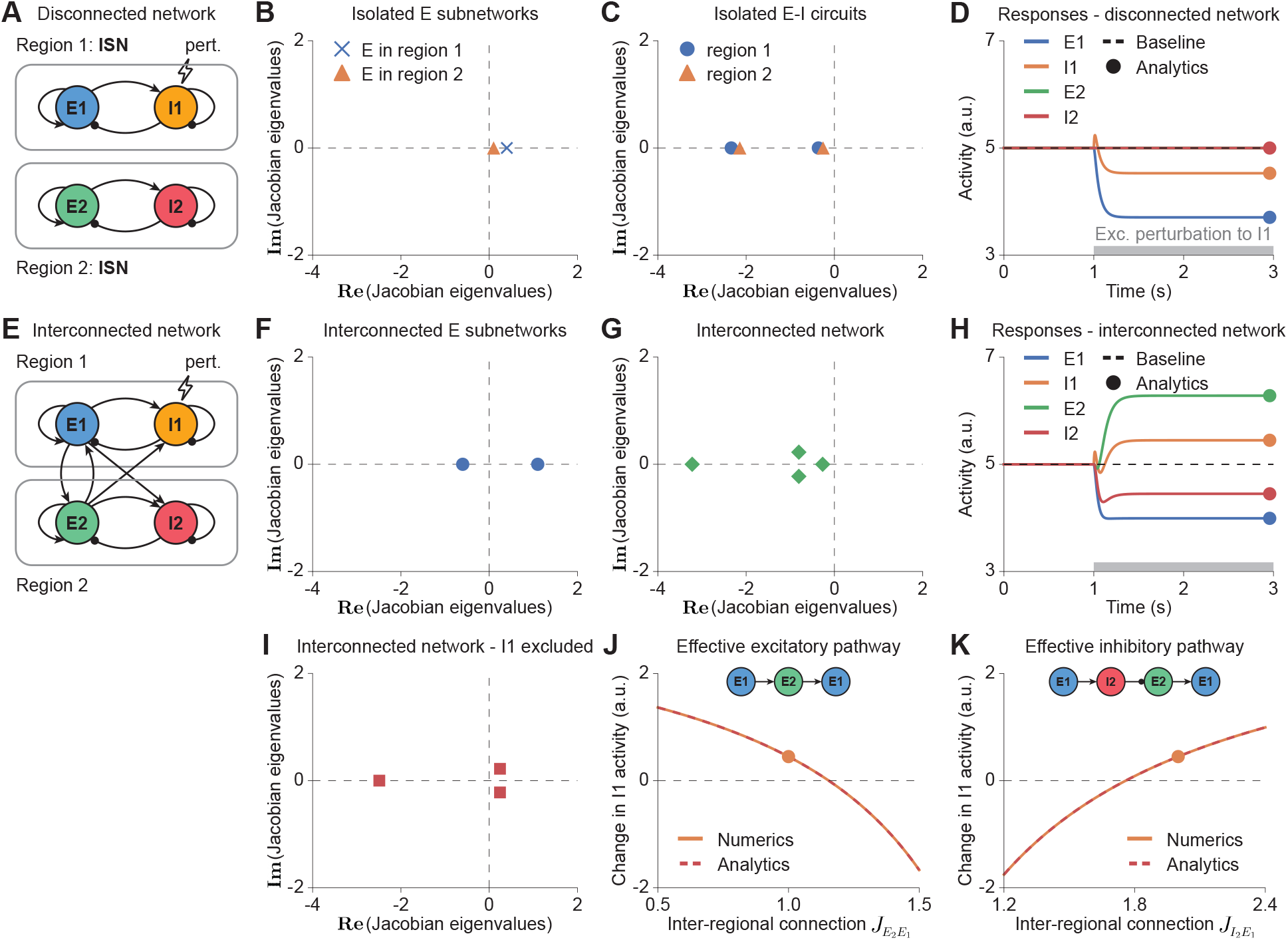
Inter-regional connections can abolish the paradoxical inhibitory response in an ISN coupled to an ISN. Panel organization, perturbation protocol, and plotting conventions follow Figure 1. **A–D**. Disconnected configuration. Both regions are ISNs in isolation, and *I*_1_ activity decreases paradoxically in response to the excitatory perturbation. **E–K**. Interconnected configuration. **E–H**. Inter-regional coupling reverses the *I*_1_ response to a nonparadoxical increase while resulting in one unstable mode in the coupled excitatory subnetwork and preserving overall network stability. **I**. The interconnected network with *I*_1_ excluded has two (an even number of) unstable eigenmodes, consistent with the nonparadoxical response in **H. J–K**. The unperturbed region (region 2) has zero (an even number of) unstable eigenmodes, as shown in **C**. For the parameters shown, increasing *J*_*E*2*E*1_, which contributes to positive inter-regional feedback, decreases the perturbation-induced change in *I*_1_ activity, whereas increasing *J*_*I*2*E*1_, which contributes to negative inter-regional feedback, increases this change.

## Discussion

Paradoxical inhibitory responses are widely regarded as a hallmark of inhibition-stabilized networks and are commonly used to infer whether a local circuit operates in an inhibition-stabilized regime. Our results show that this inference does not generally hold when the perturbed region is embedded within an interconnected network. Inter-regional coupling can generate a paradoxical response in a region that is not locally inhibition-stabilized or abolish the response in a region that is, thereby dissociating paradoxical responses from local inhibition stabilization. A paradoxical response is therefore neither necessary nor sufficient to establish local inhibition stabilization. In an interconnected network, the response is jointly determined by recurrent excitation within the perturbed region, feedback mediated through other regions, and the dynamics of the populations mediating that feedback.

By decomposing inter-regional feedback into connectivity pathways, we provide a mechanistic interpretation of this dissociation. When the subsystem comprising the other regions is stable with the perturbed region held fixed and the pathway expansion converges absolutely, pathways containing an even number of inhibitory connections provide positive feedback, whereas those containing an odd number provide negative feedback. Effective excitatory feedback can therefore arise from purely excitatory pathways or through disinhibition. These opposing contributions shape the response to perturbation. Following a positive perturbation to the local inhibitory population, local excitatory activity is initially suppressed, reducing the recurrent excitatory drive to the inhibitory population. Effective excitatory pathways reinforce this suppression and can produce a paradoxical response if the resulting loss of excitatory drive outweighs the direct perturbation. Conversely, effective inhibitory pathways counteract the suppression and can prevent a paradoxical response despite local inhibition stabilization.

The biological relevance of these effects depends on the relative strength of local and inter-regional feedback. Local recurrent connectivity is often dense, but a given region may also receive convergent long-range projections from many other regions. Consequently, even if individual long-range projections are relatively sparse or weak, their combined influence may be substantial and comparable to that of local recurrent connections. The contribution of inter-regional feedback will depend not only on the number and strength of these projections but also on the dynamics of the connected populations. Future work will be needed to determine the relative contributions of local recurrent and inter-regional connectivity.

Beyond its magnitude, the effect of inter-regional feedback depends critically on the stability of the subsystem comprising all other regions when activity in the perturbed region is held fixed. If this subsystem is unstable, with an odd number of unstable modes, the direction of the paradoxical-response inequality is reversed relative to the case in which it is stable. In biological networks, local feedback within individual regions and feedback distributed across multiple regions can both contribute to stabilization. Holding one region fixed therefore need not destabilize the remaining network, provided that these interactions do not depend critically on feedback through that region. Future experiments are needed to determine how strongly the stability of the remaining network depends on feedback from individual regions.

Experimentally determining local inhibition stabilization requires accounting for inter-regional feed-back while preserving the local operating point. In principle, closed-loop control could maintain activity in connected regions near baseline while the local inhibitory population is perturbed. This approach could approximately preserve baseline input from connected regions while limiting perturbation-induced changes in that input. If the resulting local E–I circuit remains stable, a paradoxical inhibitory response would provide strong evidence for local inhibition stabilization at the original operating point.

In our model, each brain region is represented by two populations, one excitatory and one inhibitory, each described by its mean firing rate. This population-level description makes the contributions of local and inter-regional feedback explicit, but omits several features of biological circuits, including heterogeneity among neurons within each population (Sadeh et al., 2017; Gast et al., 2024), feature-specific connectivity (Sadeh and Clopath, 2020; Chau et al., 2025; Hendricks et al., 2026), cell-type diversity (Litwin-Kumar et al., 2016; Palmigiano et al., 2020; Mahrach et al., 2020; Richter and Gjorgjieva, 2022; Waitzmann et al., 2024; Bos et al., 2025), higher-order connectivity statistics (Milo et al., 2002; Hu et al., 2013; Lin et al., 2024; Shao et al., 2025; Dahmen et al., 2026), and cell-type-specific long-range projections (Zhang et al., 2014; Shen et al., 2022). Recent theoretical work has shown that higher-order connectivity motifs can modify the relationship between inhibition stabilization and paradoxical responses even within a single region (Shao et al., 2025). Recent studies have further suggested that networks comprising multiple inhibitory cell types can operate in distinct cell-type-specific inhibition-stabilized regimes under different stimulus conditions and exhibit corresponding cell-type-specific paradoxical responses (Waitzmann et al., 2024; Cammarata et al., 2025). In networks with heterogeneous neuronal activity, some of these regimes can be detected only through patterned perturbations that vary in magnitude across neurons of the same cell type, rather than uniform perturbations applied equally to all neurons of that type (Wu et al., 2026). How these biological features modify the interplay between local and inter-regional feedback remains an open question. Future models incorporating these features will be needed to determine how local and inter-regional feedback jointly shape inhibition stabilization and paradoxical responses in more biologically realistic circuits.

More broadly, our work raises a general challenge for interpreting perturbation experiments in distributed networks. Although a perturbation may be applied to a single region, its effect can propagate through connected regions and subsequently feed back to the perturbed region. The observed response therefore reflects the dynamics of the interconnected network rather than those of the local circuit alone. Similar considerations may extend beyond paradoxical inhibitory responses to other dynamical phenomena commonly attributed to local circuits. Understanding how distributed interactions shape local computations will therefore be essential for linking perturbation experiments to the underlying circuit mechanisms.

### Numerical Simulations

Simulations were performed in Python using threshold-linear activation functions with a threshold of zero. The network dynamics were numerically integrated using the adaptive-step solver scipy.integrate.solve ivp, with a maximum step size of 1 ms. For every connectivity configuration, the baseline external input was chosen such that all four populations had a steady-state firing rate of 5 before perturbation. Because all populations had positive firing rates, their gains were equal to one. The required baseline input was consequently calculated from the steady-state equation **g**_0_ = (**I** − **J**) **r**_0_. The baseline inputs were also adjusted when assessing how inter-regional connection strength affects the inhibitory response to perturbation, maintaining the same baseline firing rates across conditions. At *t* = 1 s, a constant excitatory perturbation of amplitude 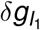 was applied to the inhibitory population in region 1 and maintained for the remainder of the simulation. All simulation parameters are listed in Tables 1 and 2.

**Table 1.** Model parameters used in Figures 1 and S1-S3.

| Symbol | Fig. 1 | Fig. S1 | Fig. S2 | Fig. S3 | Unit | Description |
| --- | --- | --- | --- | --- | --- | --- |
| $\tau_E$ | 20 | 20 | 20 | 20 | ms | time constant of E rate dynamics |
| $\tau_I$ | 20 | 20 | 20 | 20 | ms | time constant of I rate dynamics |
| $J_{E_1 E_1}$ | 0.5 | 0.3 | 0.4 | 0.3 | a.u. | connection strength from E to E in region 1 |
| $J_{I_1 E_1}$ | 0.7 | 0.5 | 1.0 | 0.5 | a.u. | connection strength from E to I in region 1 |
| $J_{E_1 I_1}$ | 0.9 | 1.2 | 0.4 | 1.0 | a.u. | connection strength from I to E in region 1 |
| $J_{I_1 I_1}$ | 0.9 | 1.3 | 0.2 | 1.1 | a.u. | connection strength from I to I in region 1 |
| $J_{E_2 E_2}$ | 0.8 | 1.2 | 3.4 | 1.7 | a.u. | connection strength from E to E in region 2 |
| $J_{I_2 E_2}$ | 1.5 | 1.3 | 3.3 | 1.1 | a.u. | connection strength from E to I in region 2 |
| $J_{E_2 I_2}$ | 0.3 | 0.9 | 3.4 | 1.0 | a.u. | connection strength from I to E in region 2 |
| $J_{I_2 I_2}$ | 1.6 | 1.0 | 1.3 | 0.9 | a.u. | connection strength from I to I in region 2 |
| $J_{E_2 E_1}$ | 0 (0.4) | 0 (0.9) | 0 (1.6) | 0 (1.3) | a.u. | connection strength from E in region 1 to E in region 2 |
| $J_{I_2 E_1}$ | 0 (1.1) | 0 (0.5) | 0 (0.2) | 0 (1.6) | a.u. | connection strength from E in region 1 to I in region 2 |
| $J_{E_1 E_2}$ | 0 (1.4) | 0 (0.9) | 0 (1.8) | 0 (0.1) | a.u. | connection strength from E in region 2 to E in region 1 |
| $J_{I_1 E_2}$ | 0 (1.9) | 0 (1.3) | 0 (1.7) | 0 (1.6) | a.u. | connection strength from E in region 2 to I in region 1 |
| $\delta g_{I_1}$ | 1 | 1 | 1 | 1 | a.u. | amplitude of the perturbation applied to I in region 1 |

Values in parentheses are used for the corresponding interconnected networks.

**Table 2.**
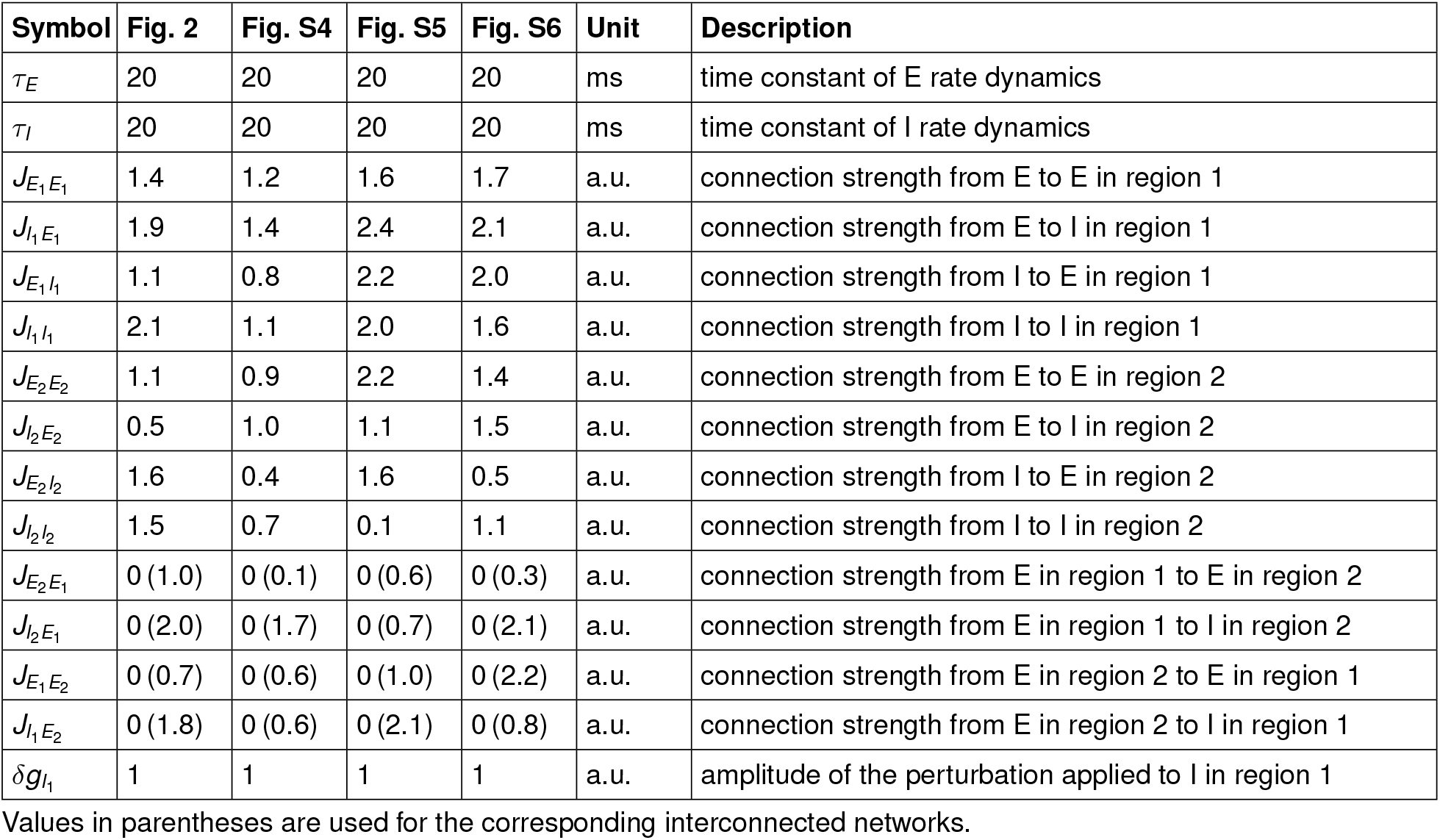
Model parameters used in Figures 2 and S4-S6.

## Data Availability

The code used for model simulations is available at https://github.com/yuekriswu/inter-regional-paradoxical-response.

## Contributions

Conceptualization, Y.K.W. and K.D.M.; methodology, Y.K.W.; formal analysis, Y.K.W.; investigation, Y.K.W.; writing–original draft, Y.K.W.; writing–review and editing, K.D.M. and Y.K.W.; funding acquisition, K.D.M.

## Acknowledgments

We thank Matthew C Rosen for commenting on the manuscript. This work is supported by the Gatsby Charitable Foundation (GAT3708), the NIH (1RF1DA056397, U19NS107613), and the Simons Foundation (SCGB 543017 to K.D.M.).

---

**Fig. S1.**
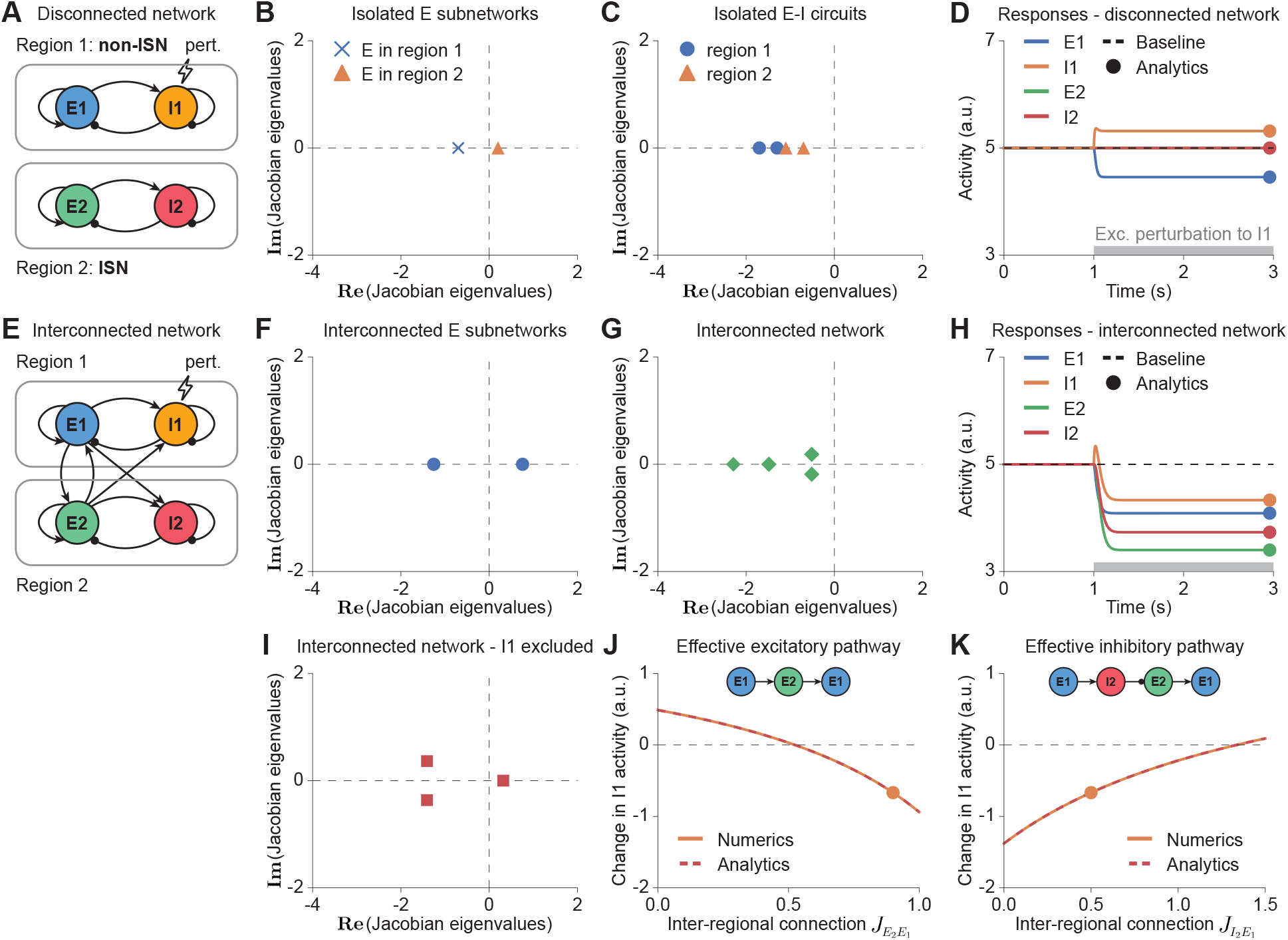
Inter-regional connections can induce a paradoxical inhibitory response in a non-ISN coupled to an ISN. Panel organization, perturbation protocol, and plotting conventions follow Figure 1. **A–D**. Disconnected configuration. In isolation, region 1 is a non-ISN, whereas region 2 is an ISN. *I*_1_ activity increases nonparadoxically in response to the excitatory perturbation. **E–K**. Interconnected configuration. **E–H**. Inter-regional coupling reverses the *I*_1_ response to a paradoxical decrease while resulting in one unstable mode in the coupled excitatory subnetwork and preserving overall network stability. **I**. The interconnected network with *I*_1_ excluded has an odd number of unstable eigenmodes (one in this case), consistent with the paradoxical response in **H. J–K**. The unperturbed region (region 2) has zero (an even number of) unstable eigenmodes, as shown in **C**. Increasing *J*_*E*2*E*1_, which contributes to positive inter-regional feedback, decreases the perturbation-induced change in *I*_1_ activity, whereas increasing *J*_*I*2*E*1_, which contributes to negative inter-regional feedback, increases this change.

**Fig. S2.**
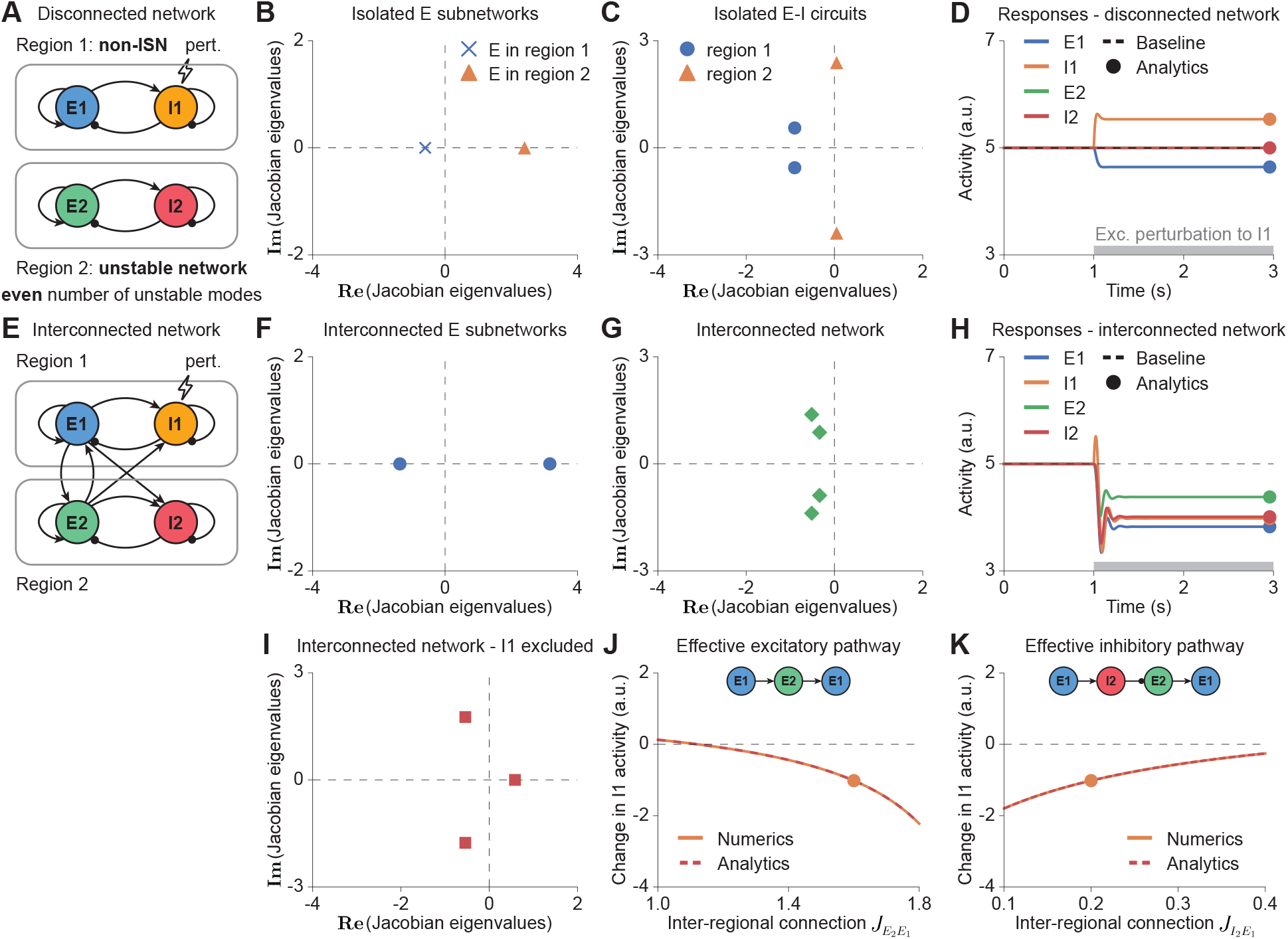
Inter-regional connections can induce a paradoxical inhibitory response in a non-ISN coupled to an unstable network with an even number of unstable eigenmodes. Panel organization, perturbation protocol, and plotting conventions follow Figure 1. **A–D**. Disconnected configuration. In isolation, region 1 is a non-ISN, whereas region 2 is an unstable network with two (an even number of) unstable eigenmodes. *I*_1_ activity increases nonparadoxically in response to the excitatory perturbation. **E–K**. Interconnected configuration. **E–H**. Inter-regional coupling reverses the *I*_1_ response to a paradoxical decrease while resulting in one unstable mode in the coupled excitatory subnetwork and stabilizing the full interconnected network. **I**. The interconnected network with *I*_1_ excluded has an odd number of unstable eigenmodes (one in this case), consistent with the paradoxical response in **H. J–K**. The unperturbed region (region 2) has two (an even number of) unstable eigenmodes, as shown in **C**. Increasing *J*_*E*2*E*1_, which contributes to positive inter-regional feedback, decreases the perturbation-induced change in *I*_1_ activity, whereas increasing *J*_*I*2*E*1_, which contributes to negative inter-regional feedback, increases this change.

**Fig. S3.**
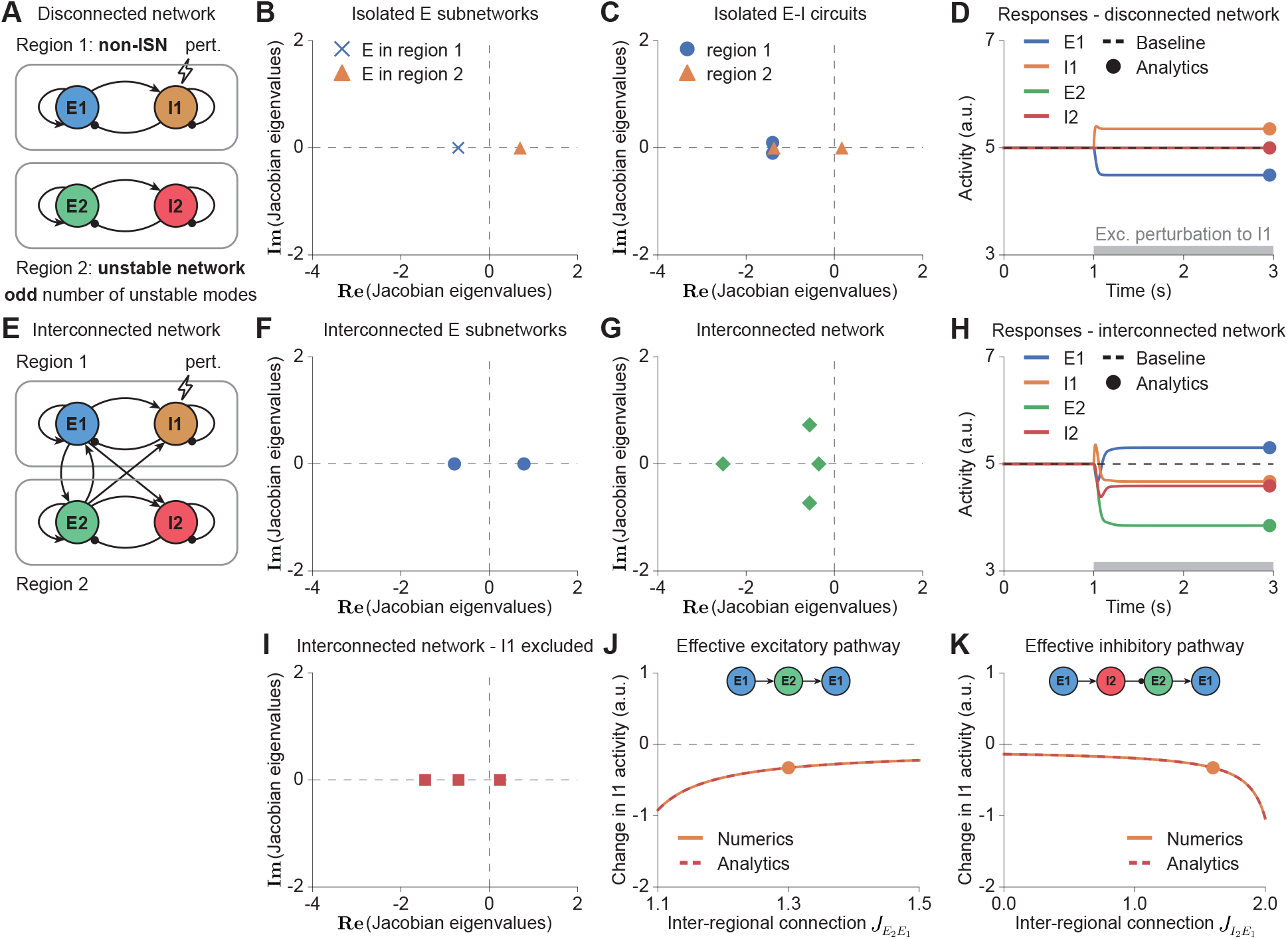
Inter-regional connections can induce a paradoxical inhibitory response in a non-ISN coupled to an unstable network with an odd number of unstable eigenmodes. Panel organization, perturbation protocol, and plotting conventions follow Figure 1. **A–D**. Disconnected configuration. In isolation, region 1 is a non-ISN, whereas region 2 is an unstable network with a single unstable eigenmode and thus an odd number of unstable eigenmodes. *I*_1_ activity increases nonparadoxically in response to the excitatory perturbation. **E–K**. Interconnected configuration. **E–H**. Inter-regional coupling reverses the *I*_1_ response to a paradoxical decrease while resulting in one unstable mode in the coupled excitatory subnetwork and stabilizing the full interconnected network. **I**. The interconnected network with *I*_1_ excluded has an odd number of unstable eigenmodes (one in this case), consistent with the paradoxical response in **H. J–K**. The unperturbed region (region 2) has an odd number of unstable eigenmodes (one in this case), as shown in **C**. Unlike in Figures 1, S1, and S2, increasing *J*_*E*2*E*1_, which is involved in the effective excitatory pathway *E*_1_ → *E*_2_ → *E*_1_, increases the perturbation-induced change in *I*_1_ activity, whereas increasing *J*_*I*2*E*1_, which is involved in the effective inhibitory pathway *E*_1_ → *I*_2_ → *E*_2_ → *E*_1_, decreases this change.

**Fig. S4.**
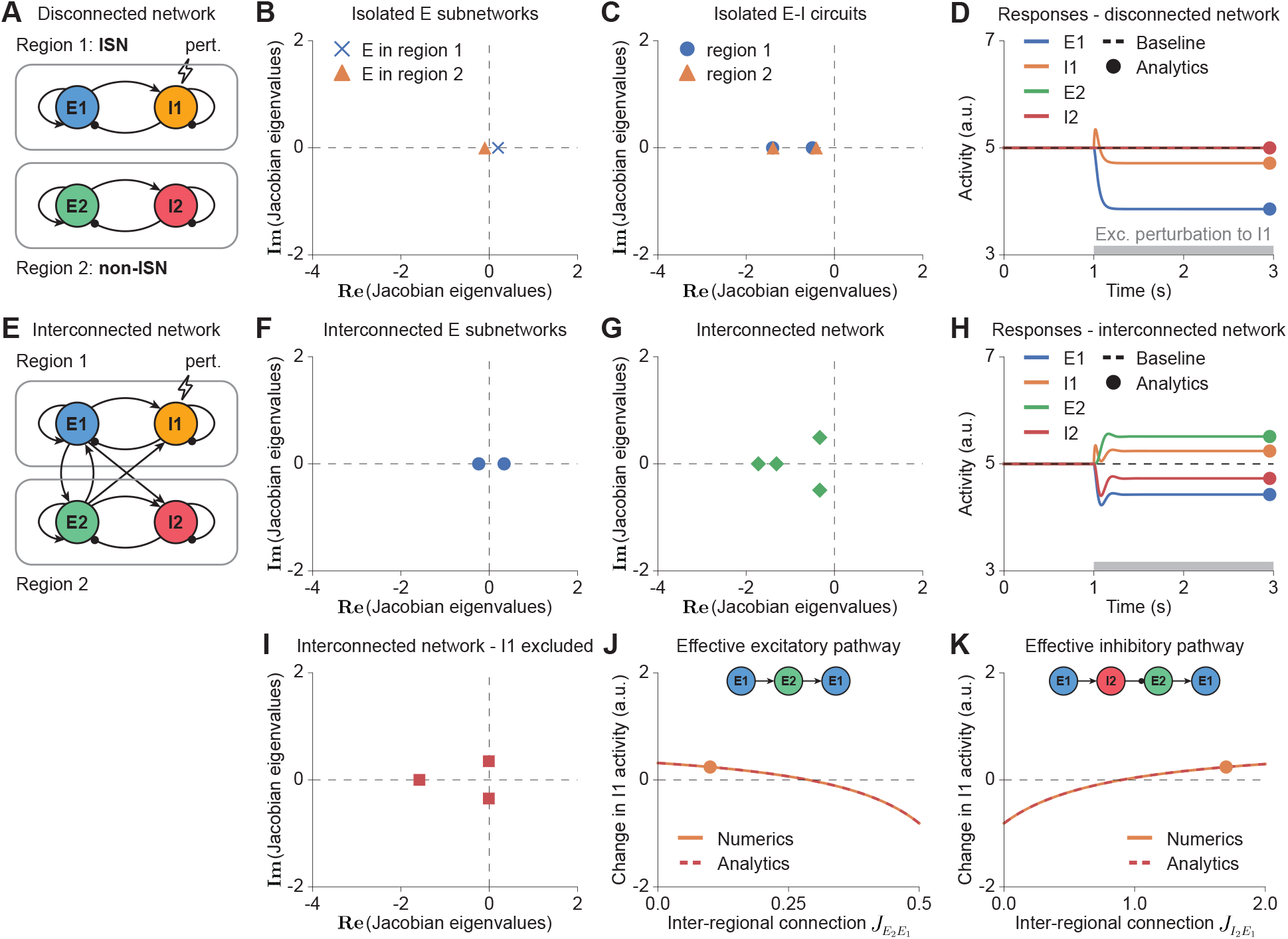
Inter-regional connections can abolish the paradoxical inhibitory response in an ISN coupled to a non-ISN. Panel organization, perturbation protocol, and plotting conventions follow Figure 1. **A–D**. Disconnected configuration. In isolation, region 1 is an ISN, whereas region 2 is a non-ISN. *I*_1_ activity decreases paradoxically in response to the excitatory perturbation. **E–K**. Interconnected configuration. **E–H**. Inter-regional coupling reverses the *I*_1_ response to a nonparadoxical increase while resulting in one unstable mode in the coupled excitatory subnetwork and preserving overall network stability. **I**. The interconnected network with *I*_1_ excluded has zero (an even number of) unstable eigenmodes, consistent with the nonparadoxical response in **H. J–K**. The unperturbed region (region 2) has zero (an even number of) unstable eigenmodes, as shown in **C**. For the parameters shown, increasing *J*_*E*2*E*1_, which contributes to positive inter-regional feedback, decreases the perturbation-induced change in *I*_1_ activity, whereas increasing *J*_*I*2*E*1_, which contributes to negative inter-regional feedback, increases this change.

**Fig. S5.**
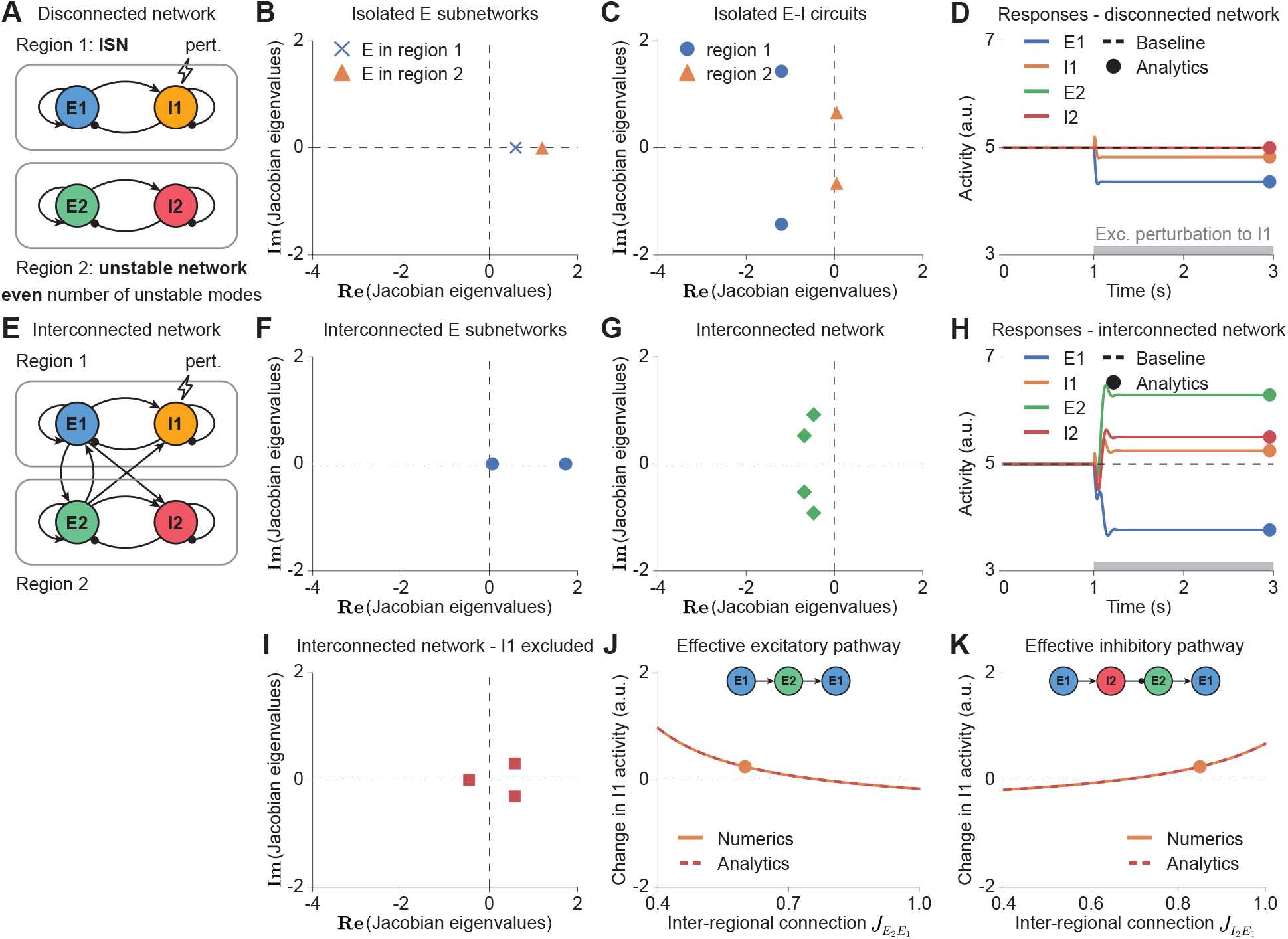
Inter-regional connections can abolish the paradoxical inhibitory response in an ISN coupled to an unstable network with an even number of unstable eigenmodes. Panel organization, perturbation protocol, and plotting conventions follow Figure 1. **A–D**. Disconnected configuration. In isolation, region 1 is an ISN, whereas region 2 is an unstable network with two (an even number of) unstable eigenmodes. *I*_1_ activity decreases paradoxically in response to the excitatory perturbation. **E–K**. Interconnected configuration. **E–H**. Inter-regional coupling reverses the *I*_1_ response to a nonparadoxical increase while resulting in two unstable modes in the coupled excitatory subnetwork and stabilizing the full interconnected network. **I**. The interconnected network with *I*_1_ excluded has two (an even number of) unstable eigenmodes, consistent with the nonparadoxical response in **H. J–K**. The unperturbed region (region 2) has two (an even number of) unstable eigenmodes, as shown in **C**. For the parameters shown, increasing *J*_*E*2*E*1_, which contributes to positive inter-regional feedback, decreases the perturbation-induced change in *I*_1_ activity, whereas increasing *J*_*I*2*E*1_, which contributes to negative inter-regional feedback, increases this change.

**Fig. S6.**
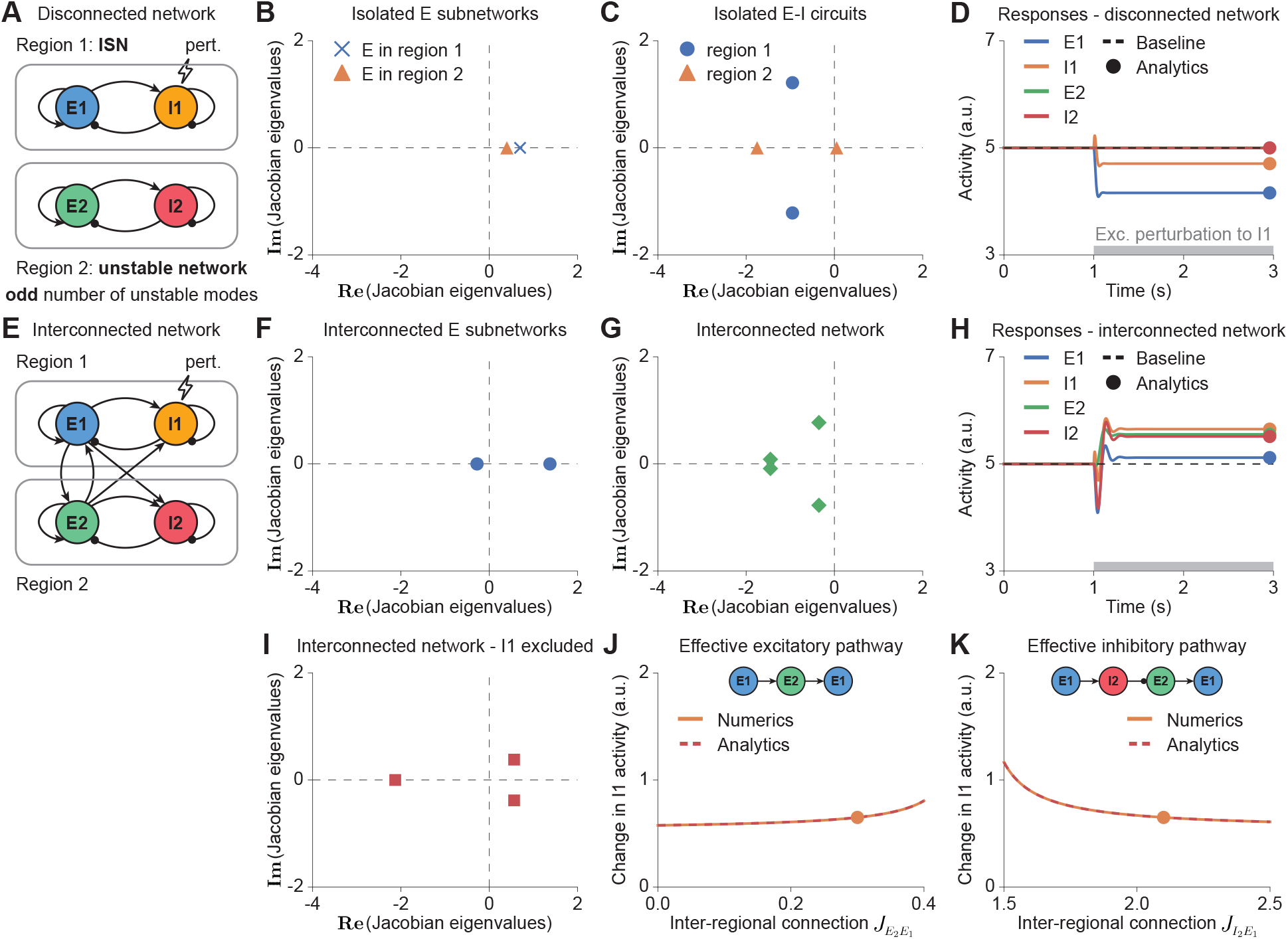
Inter-regional connections can abolish the paradoxical inhibitory response in an ISN coupled to an unstable network with an odd number of unstable eigenmodes. Panel organization, perturbation protocol, and plotting conventions follow Figure 1. **A–D**. Disconnected configuration. In isolation, region 1 is an ISN, whereas region 2 is an unstable network with a single unstable eigenmode and thus an odd number of unstable eigenmodes. *I*_1_ activity decreases paradoxically in response to the excitatory perturbation. **E–K**. Interconnected configuration. **E–H**. Inter-regional coupling reverses the *I*_1_ response to a nonparadoxical increase while resulting in one unstable mode in the coupled excitatory subnetwork and stabilizing the full interconnected network. **I**. The interconnected network with *I*_1_ excluded has two (an even number of) unstable eigenmodes, consistent with the nonparadoxical response in **H. J–K**. The unperturbed region (region 2) has an odd number of unstable eigenmodes (one in this case), as shown in **C**. For the parameters shown, unlike in Figures 2, S4, and S5, increasing *J*_*E*2*E*1_, which is involved in the effective excitatory pathway *E*_1_ → *E*_2_ → *E*_1_, increases the perturbation-induced change in *I*_1_ activity, whereas increasing *J*_*I*2*E*1_, which is involved in the effective inhibitory pathway *E*_1_ → *I*_2_ → *E*_2_ → *E*_1_, decreases this change.

